# *De novo* assembly of a young Drosophila Y chromosome using Single-Molecule sequencing and Chromatin Conformation capture

**DOI:** 10.1101/324673

**Authors:** Shivani Mahajan, Kevin Wei, Matthew Nalley, Lauren Giblisco, Doris Bachtrog

## Abstract

While short-read sequencing technology has resulted in a sharp increase in the number of species with genome assemblies, these assemblies are typically highly fragmented. Repeats pose the largest challenge for reference genome assembly, and pericentromeric regions and the repeat-rich Y chromosome are typically ignored from sequencing projects. Here, we assemble the genome of *Drosophila miranda* using long reads for contig formation, chromatin interaction maps for scaffolding and short reads, optical mapping and BAC clone sequencing for consensus validation. Our assembly recovers entire chromosomes and contains large fractions of repetitive DNA, including ^~^41.5 Mb of pericentromeric and telomeric regions, and >100Mb of the recently formed highly repetitive neo-Y chromosome. While Y chromosome evolution is typically characterized by global sequence loss and shrinkage, the neo-Y increased in size by almost 3-fold, due to the accumulation of repetitive sequences. Our high-quality assembly allows us to reconstruct the chromosomal events that have led to the unusual sex chromosome karyotype in *D. miranda*, including the independent *de novo* formation of a pair of sex chromosomes at two distinct time points, or the reversion of a former Y chromosome to an autosome.

## Introduction

Sex chromosomes are derived from ordinary autosomes, yet old X and Y chromosomes contain a vastly different gene repertoire [1]. In particular, X chromosomes resemble the autosome from which they were derived, with only few changes to their gene content [2]. In contrast, Y chromosomes dramatically remodel their genomic architecture. Y evolution is characterized by massive gene decay, with the vast majority of the genes originally present on the Y disappearing, and Y degeneration is often accompanied by the acquisition of repetitive DNA [3]; old Y chromosomes typically have shrunk dramatically in size and contain only few unique genes but vast amounts of repeats.

The decrease in sequencing cost and increased sophistication of assembly algorithms for short-read platforms has resulted in a sharp increase in the number of species with genome assemblies. Indeed, X chromosomes have been characterized and sequenced in many species. However, assemblies based on short-read technology are highly fragmented, with many gaps, ambiguities, and errors remaining; this is especially true for repeat-rich regions, such as centromeres, telomeres, or the Y chromosome [4–6]. Thus, most sequencing projects have ignored the Y chromosome. Labor intensive sequencing of Y chromosomes in a few mammal species has revealed a surprisingly dynamic history of Y chromosome evolution, with meiotic conflicts driving gene acquisition on the mouse Y chromosome [7], or gene conversion within palindromes retarding Y degeneration in primates [8]. However, all current Y assemblies are based on tedious re-sequencing of BAC clones and available only for a handful of species [9–11], and the repeat-rich nature of Y chromosomes has hampered their evolutionary studies in most organisms.

Here we present a near-finished reference genome for *Drosophila miranda*, including its Y chromosome, using a combination of long-read single-molecule sequencing, high-fidelity short-read sequencing, optical mapping, BAC clones sequencing, and Hi-C-based chromatin interaction maps. *D. miranda* has become a model system for studying the molecular and evolutionary processes driving sex chromosome differentiation, due to its recently evolved neo-sex chromosome system (see Figure 1). In particular, chromosomal fusions within *D. miranda* have resulted in the recent sex-linkage of former autosomes at two independent time points (Figure 1D), and these new sex chromosomes are at different stages in their transition to differentiated sex chromosomes. Specifically, chromosome XR and YD became sex-linked about 15 million years ago [12], and the neo-X and neo-Y became sex chromosomes only about 1.5 million years ago [13]. These former autosomes are in the process of evolving the stereotypical properties of ancestral sex chromosomes [14,15]. Intriguingly, the ancestral Y chromosome (Y_anc_) in this species group became fused to an autosome, probably around the same time when XR and YD formed, and lost some of the characteristics of an ancient Y chromosome [16–18]. Thus, *D. miranda* allows the investigation of the functional and evolutionary changes occurring on differentiating sex chromosomes, and their reversal.

**Figure 1.**
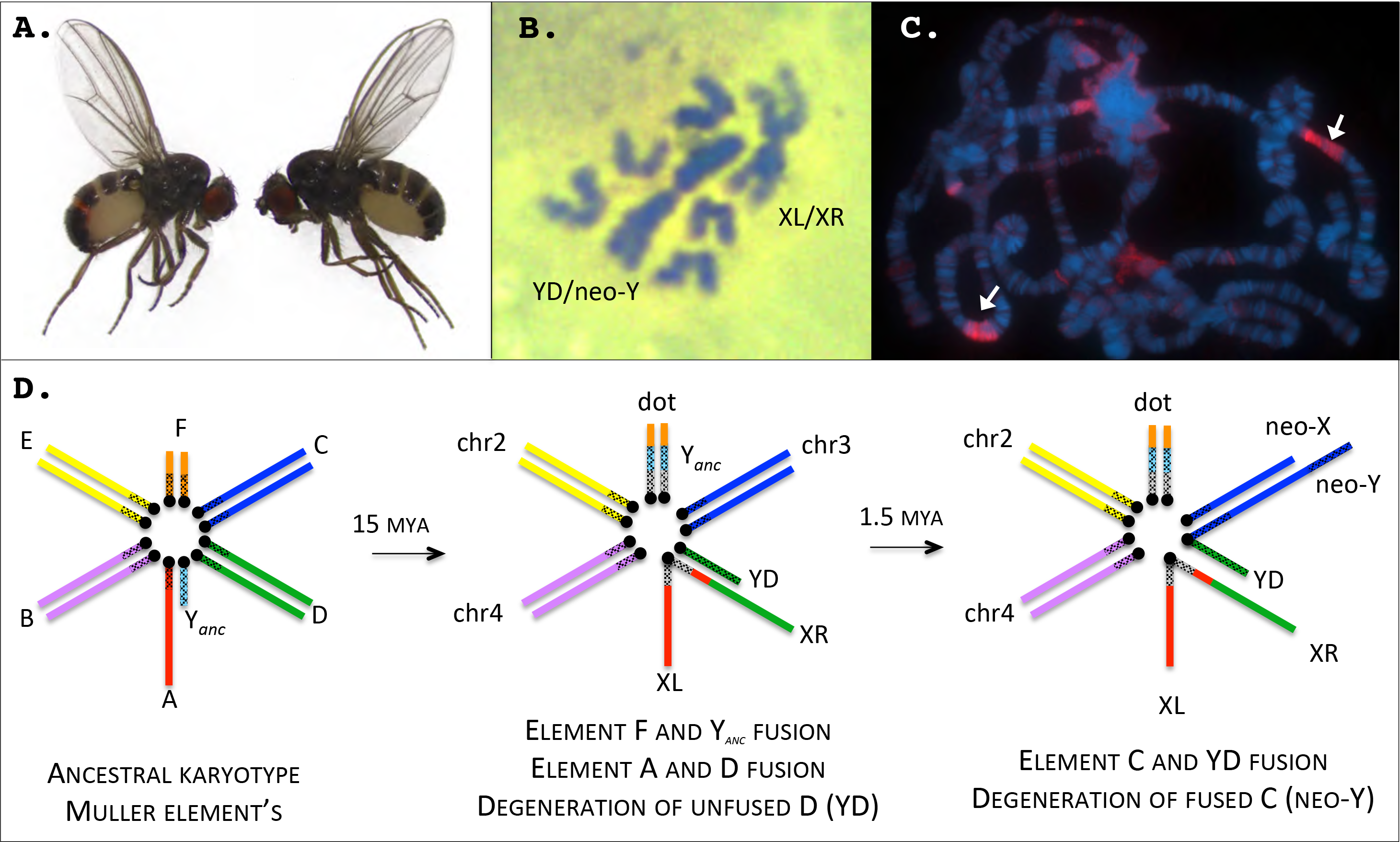
*Drosophila miranda* is a model species to study sex chromosome evolution. **A.** Male (left) and female (right) *D. miranda*. **B.** Mitotic chromosome squashes of male *D. miranda*. Both the ancestral X (XL/XR) and the Y chromosome (YD/neo-Y) show large blocks of dark staining (Giemsa), indicative of heterochromatin. The acrocentric rods are the neo-X, and chromosome 2 and 4. **C.** Polytene chromosomes of a female *D. miranda* stained for *HP1* (heterochromatin protein 1). Note the large blocks of heterochromatin (arrows) on chromosome 2 and 4. **D.** Karyotype evolution in *D. miranda*. Chromosomal fusions between the sex chromosomes and autosomes have resulted in both the reversal of an ancestral Y chromosome (Y_anc_) to an autosome, as well as the independent *de novo* formation of new sex chromosomes from autosomes at two distinct evolutionary time points (XR and YD were formed about 15MY ago, and the neo-X and neo-Y originated about 1.5 MY ago). Genome analysis allows us to reconstruct the temporal dynamics and molecular processes involved in sex chromosome evolution in this species.

The most recent assembly of *D. miranda* was generated via short-read Illumina sequencing and is highly fragmented. In particular, the genome was in 47,035 scaffolds, with a scaffold N50 of 5,007 bp and a total assembled genome size of 112 Mb (a female-only assembly resulted in 22,259 scaffolds, with an N50 of 13,773 bp and an assembled size of 125 Mb). The high amount of sequence similarity between the neo-sex chromosomes (98.5% identical at the nucleotide level), yet high repeat content of the neo-Y (over 50% of its DNA is derived from repeats) posed a particular challenge to its assembly using short reads, and our initial attempts to assemble the neo-Y resulted in a chimeric, highly fragmented and incomplete assembly, consisting of 36,282 (often chimeric) scaffolds, and a scaffold N50 of only 715 bp [19]. Thus, our previous analysis of neo-Y chromosome gene content evolution was instead based on mapping male reads to the neo-X assembly and identifying male-specific SNPs [19], or trying to reconstruct neo-Y transcripts using both male and female genome and transcriptome data [20]. This indirect approach, however, only allows the investigation of conserved regions on the neo-sex chromosome that differ by simple SNPs or short indels within genes. Here, we assemble the genome of *D. miranda* using long reads for contig formation, short reads for consensus validation, and scaffolding by chromatin interaction mapping, and we verify our assembly using optical maps and BAC clone sequencing. Our assembly covers large fractions of repetitive DNA, with entire chromosomes being in a single scaffold, including their centromeres, and we recover over >100Mb of the recently formed neo-Y chromosome. Our new assembly strategy achieves superior continuity and accuracy, and provides a new standard reference for the investigation of repetitive sequences and Y chromosome evolution in this species.

## Results & Discussion

### *De novo* assembly of a *D. miranda* reference genome

We sequenced adult male *D. miranda* (from the inbred strain MSH22), using a combination of different technologies: single-molecule real-time sequencing (PacBio), paired-end short-read sequencing (Illumina HiSeq), optical mapping (using BioNano), shot-gun BAC clones sequencing (Illumina HiSeq) and chromatin conformation capture (Hi-C; see **Table S1**).

Assembly of these complementary data types proceeded in a stepwise fashion (Figure 2A), similar to a recent approach [21], to produce progressively improved assemblies (Table 1). Briefly, we produced two initial assemblies of the PacBio data alone using the *Falcon* [22] and *Canu* [23] assembler, and double-merged the resulting assemblies with *Quickmerge* [24]. The resulting hybrid assembly had a contig NG50 (the minimum length of contigs accounting for half of the haploid genome size) of 5.2 Mb in 271 scaffolds. PacBio contigs were separated into X-linked and autosomal contigs versus Y-linked contigs based on genomic coverage patterns of mapped male-and female Illumina reads (**Figure S1**), to avoid cross-mapping of short read Hi-C data, and clustered into chromosome-scale scaffolds using Hi-C data (Figure 2B, **Figure S2, S3**). Mapping of Illumina reads also allowed us to identify and remove contigs that resulted from uncollapsed haplotypes (**Figure S4**). X-linked and autosomal contigs were scaffolded with female Hi-C libraries, while Y-linked contigs were clustered using Y-mapping reads from male Hi-C libraries (**Figure S2**). Visual inspection of contact probability maps allowed us to identify a few mis-assemblies, which were manually corrected followed by re-scaffolding (**Table S2**). To assess quality, the resulting assembly was validated via statistical methods and short read Illumina mapping (**Table S3**), and comparison to optical mapping data (**Table S4** & **Figure S5**), sequenced BAC clones from the MSH22 strain (**Table S5**, **S6**, Figure 2D and **Figure S6**) and previous assemblies (*D. miranda* D.mir1.0 [19] Figure 2C and **Figure S7**; *D. pseudoobscura;* **Figure S8**).

**Figure 2.**
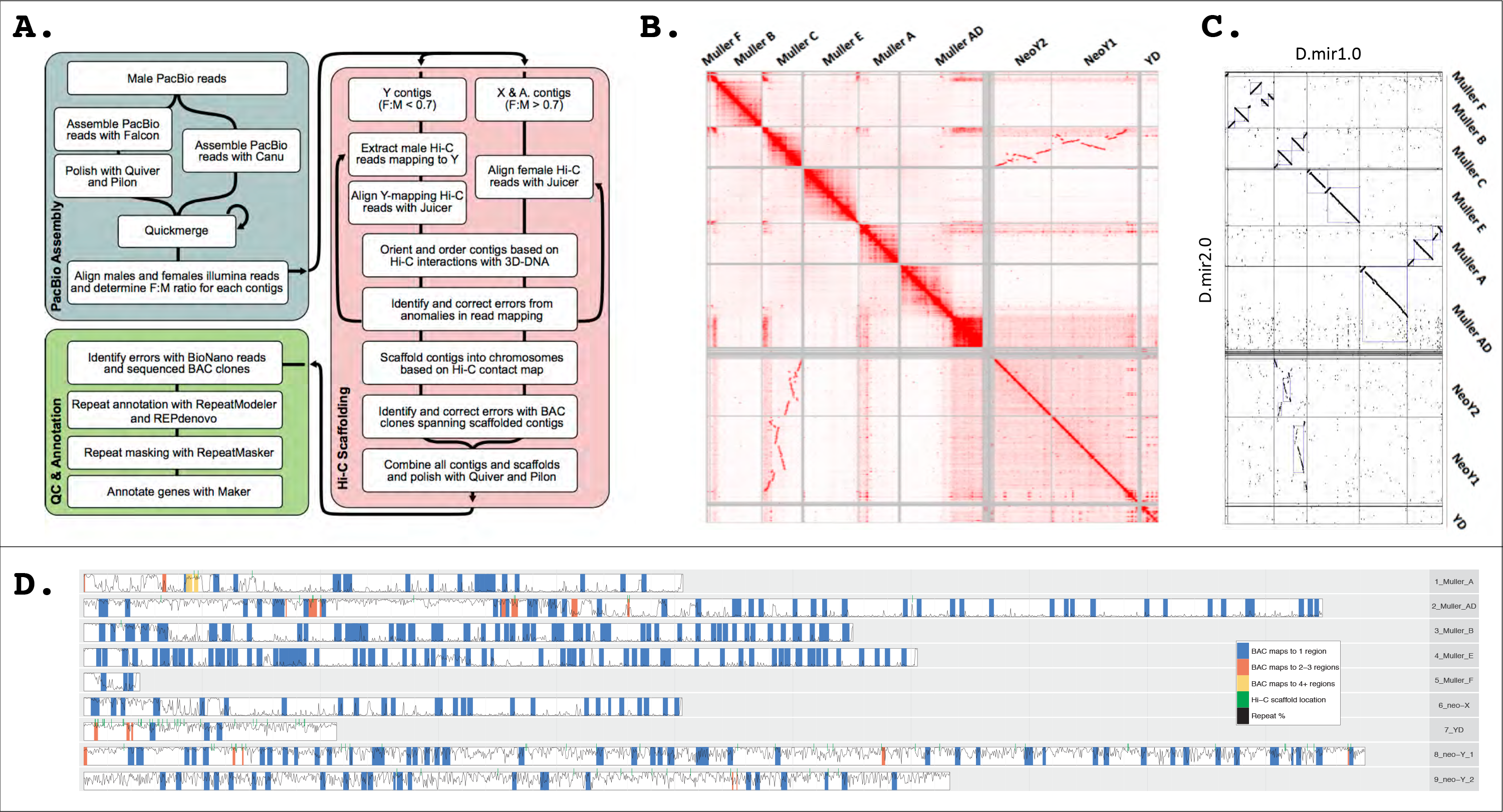
Assembly and validation of *D. miranda* genome. **A.** Overview of assembly pipeline. The steps include assembly of male PacBio reads, followed by scaffolding using Hi-C, and extensive QC using BioNano reads and BAC clone sequencing, followed by gene and repeat annotation. **B.** Hi-C linkage density map. Note that the Y-linked contigs were scaffolded separately from X-linked and autosomal contigs. Unlinked regions with many contacts indicate repetitive regions. **C.** Comparison of current (Dmir2.0) vs. old (Dmir1.0) *D. miranda* assembly. Note that the Y/neo-Y was not assembled in Dmir1.0, and the dot plot indicates homology between our neo-Y assembly, and the neo-X. Other repeat-rich regions, such as the large pericentromeric block on AD, are also missing from D.mir1.0. **D.** BAC clone mapping for assembly verification. 361 sequenced BAC clones (97%) map contiguously and uniquely to our genome assembly.

**Table 1.** Assembly statistics.

| Assembly | Contigs +<br>Scaffolds | Scaffolds | Unplaced<br>Contigs | N50 (bp) | Assembly Size<br>(Mb) | Assembly in<br>Scaffolds (%) |
| --- | --- | --- | --- | --- | --- | --- |
| PacBio Falcon | 625 | NA | 625 | 2,242,328 | 273 | NA |
| PacBio Canu | 521 | NA | 521 | 3,884,273 | 296 | NA |
| Quickmerged | 271 | NA | 271 | 5,177,776 | 295 | NA |
| PacBio+Hi-C | 103 | 14 | 89 | 37,186,217 | 289 | 96.5 |
| D.mir1.0 (female only, stitched<br>with *D.pseudoobscura*) | 13,773 | 6 | 530 | 28,826,359 | 140 | 97.9 |
| D.mir1.0 (not stitched with<br>*D. pseudoobscura*) | 47,035 | NA | NA | 5,007 | 112 | NA |
| D.mir2.0; X-linked and<br>autosomal scaffolds | 41 | 6 | 35 | 32,539,883 | 177 | 97.1 |
| D.mir2.0; Y-linked scaffolds | 62 | 8 | 54 | 36,637,378 | 111 | 95.7 |
| D.mir2.0 | 103 | 14 | 89 | 35,263,102 | 287 | 96.6 |

To maximize accuracy of the final reference assembly, errors were manually curated before final gap filling and polishing (**Table S2**). Our final assembly, D.mir2.0, totaled 287Mb of sequence with a scaffold NG50 of 35.3 Mb (Table 1). D.mir2.0 comprises just 103 scaffolds and 120 gaps (**Table S7**), and the three autosomes, the three X chromosomes and the Y of *D. miranda* are all mostly covered by a single scaffold (Figure 3). The unplaced scaffolds are relatively small (median size 37.3 kb) and highly repeat rich (median repeat content 94.7%), and coverage suggests that most are derived from the Y chromosome. In contrast, the previous assembly D.mir1.0 consisted of 47,035 scaffolds [19]. We used two approaches, REPdenovo [25] and RepeatModeler [26] to annotate repeats in the *D. miranda* genome, and Maker [27] to annotate genes (Figure 3). We identified a total of 17,745 genes, and 43.7% of the genome was annotated as repeats. BUSCO assessments [28] support that our genome assembly and annotation are highly complete (**Table S8**).

**Figure 3.**
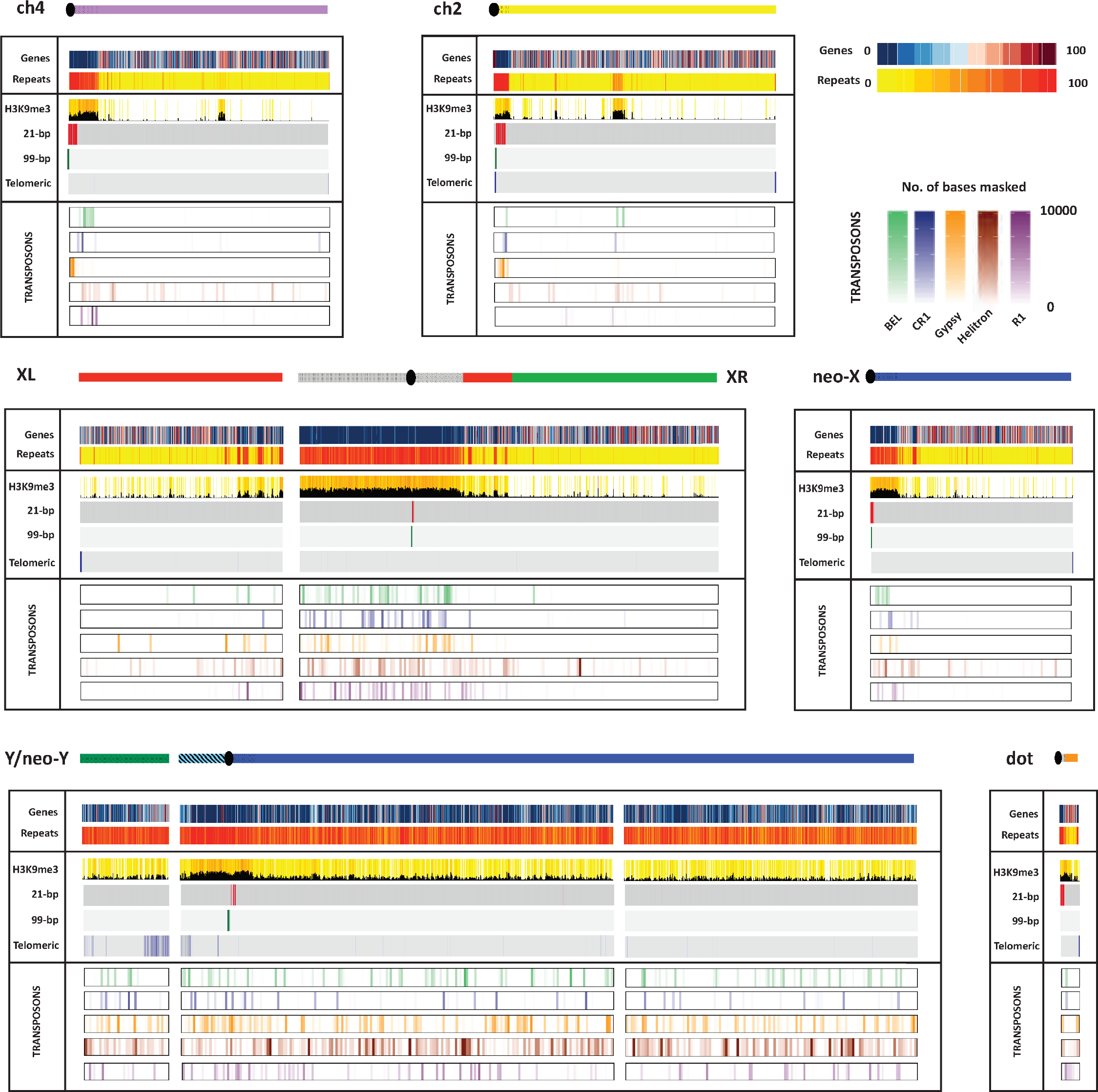
Gene and repeat content of *D. miranda* genome assembly. Shown is the gene content, repeat content, H3K9me3 enrichment and density of the most abundant satellites (21-bp repeat and 99-bp repeat), the telomeric transposable elements, and the most abundant transposons across *the D. miranda* genome assembly. A cartoon of the chromosomes is drawn with color indicating the Muller element (see Figure 1D), and the shaded regions are heterochromatic.

### Assembly benchmarking and comparison to reference

The previous *D. miranda* reference assembly (D.mir1.0) was generated from paired-end short reads using the *SOAPdenovo* assembler, and cross-species scaffold alignments to *D. pseudoobscura* [19]. Paired-end read sequences used to create the D.mir1.0 reference assembly were aligned to our D.mir2.0 assembly for a reference-free measure of structural correctness. These alignments confirmed that our current assembly is a dramatic improvement over D.mir1.0 (**Table S3**), with fewer putative translocations (36 versus 17,764), deletions (229 versus 6075) and duplications (8 versus 1703). The initial *D. miranda* genome was scaffolded using *D. pseudoobscura*, and genome-wide alignments between our current *D. miranda* assembly and D.mir1.0 reveals dozens of inversions that were introduced by the scaffolding (Figure 2C; **Figure S7**).

We independently assess the quality and large-scale structural continuity of our assembly by comparing it to sequenced BAC clones and optimal mapping data. In total, we shotgun sequenced 383 randomly selected BAC clones from a *D. miranda* male BAC clone library [29], which should cover roughly 1/4 of the *D. miranda* genome. 372 BAC clones passed our sequence coverage filter, and could be aligned to our *D. miranda* genome; of those, 361 (i.e. 97%) contiguously map to a unique position in the genome (Figure 2D; **Table S5, S6; Figure S6**). Only 11 BAC clones map to 2 or 3 (typically highly repetitive) genomic locations (Figure 2D), and could represent assembly mistakes or recombinant BAC clones. Similarly, most of our genome is covered by optical mapping data (**Figure S5 & Table S4**). Thus, continuous and unique mapping of most BAC clones and coverage by optical reads supports that our genome assembly is of high quality.

### Assembly of highly repeat-rich regions

Our high-quality assembly contains large amounts of highly repetitive regions, including telomeres, pericentromeric regions and putative centromeric repeats, as well as the repeat-rich Y chromosome. Overall, ^~^126 Mb of the assembled 287 Mb *D. miranda* genome are repetitive, and we assembled ^~^41 Mb of pericentromeric and centromeric repeats and telomeres (**Table S7**). In some cases, we assemble through the entire centromere and recover telomeric repeats at the end of a chromosome arm (see below). In contrast, the previous *D. miranda* assembly based on only Illumina reads recovered less than 0.5 Mb of pericentromeric DNA (**Table S7, Figure S7**), and even the highly curated *D. melanogaster* genome assembly [30] entirely lacks centromeric sequence (**Figure S9**). In addition, we assembled 110.5 Mb of Y-linked sequence, with 101.5 Mb contained within a single scaffold (Figure 3). Our assembly allows us to recover structural variants, including gene duplications and tandem repeats, most of which were collapsed and missed in our previous assembly (**Figure S10**).

### Recovery of chromosome ends and identification of putative centromeric DNA sequences

In Drosophila, telomeres are maintained by the occasional transposition of specific non-LTR retrotransposons (i.e. the HeT-A, TAHRE and TART elements) to chromosome ends [31,32], and hybridization studies have suggested about two telomere repeats per chromosome end in *D. miranda* [33]. Indeed, for almost all chromosome arms (Muller A, B, C, both ends of E, F, neo-Y and YD), we properly identified the ends of chromosomes based on the presence of telomeric transposable elements (see Figure 4A, B).

**Figure 4.**
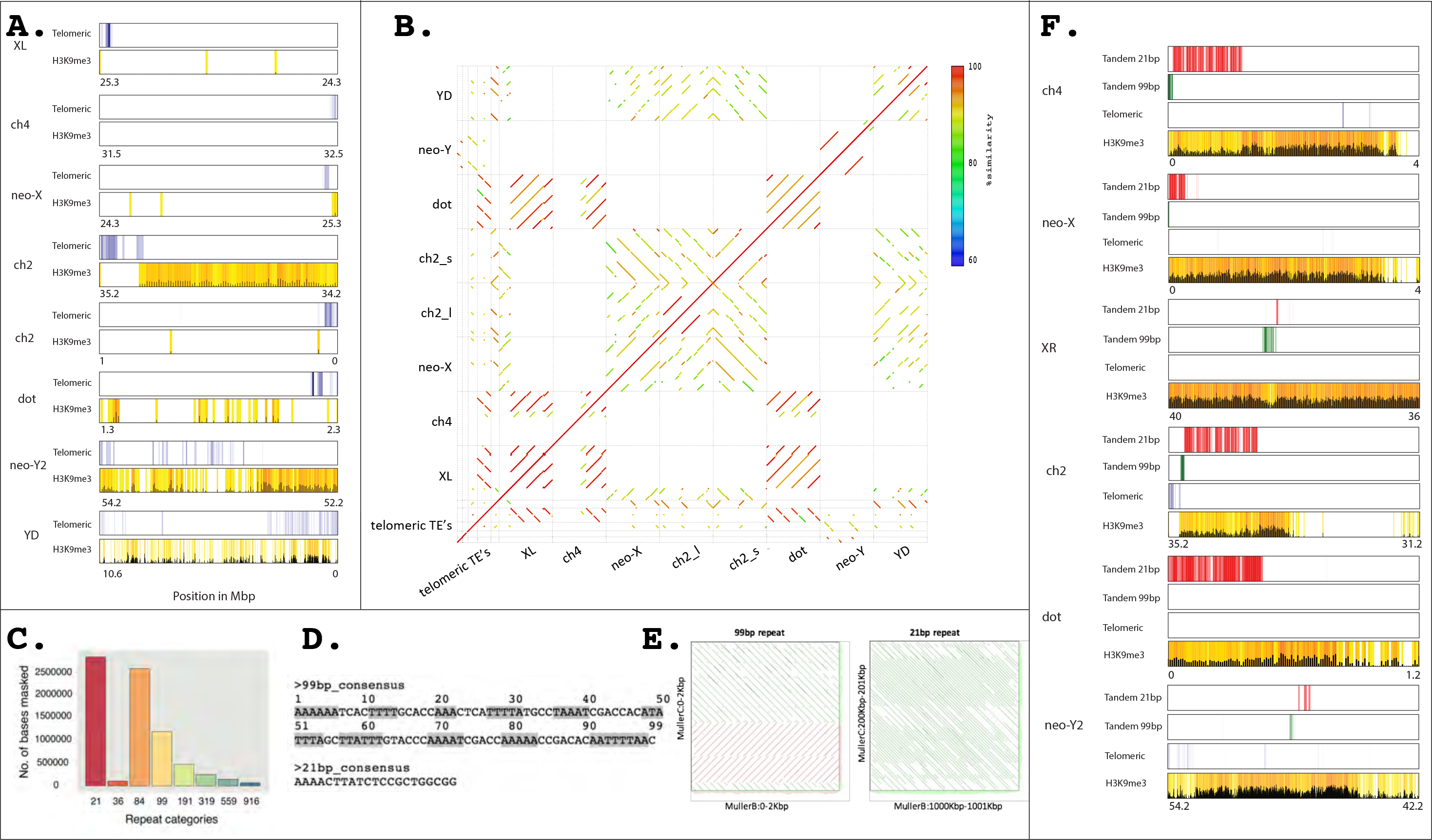
Recovery of telomeres and identification of putative centromere repeats for each chromosome. **A.** Presence of telomere repeats at or near the ends of most chromosome arms. Shown is enrichment of telomere repeats and H3K9me3 marks in a sliding window. **B.** Alignment of chromosome ends and telomere repeats. **C.** Histogram of most abundant satellite in *D. miranda* genome. Repeat categories refer to the size of the repeat unit. Note that the 84-mer is a higher-order variant of four units of the 21-mer. **D.** Consensus sequence of 21-bp and 99-bp repeat. E. Comparison of the centromeric repeat from different chromosomes. **F.** Location of putative centromere repeats in pericentromeric regions, and H3K9me3 enrichment. H3K9me3 enrichment is reduced at the putative centromeric repeat (**Figure S13**). Note that for the acrocentric chromosome 2, we recover the entire centromere, including the telomere.

Centromere sequences show little conservation between closely related species, but have a common organization in most animals and plants [34–36]. In particular, centromeres typically comprise megabase-scale arrays of tandem repeats embedded in heterochromatin, but are notoriously difficult to recover in genome assemblies. In several instances, we sequenced several megabases into the highly repetitive pericentromeric region (Figure 3, 4; **Figure S11**), and for one chromosome (Muller element E), we assembled the entire chromosome (based on the recovery of telomeric sequences on both chromosome ends), including its centromere.

We used Tandem Repeat Finder (TRF)[37] to identify satellite repeats, and plotted their occurrence along the genome (Figure 3, 4C, **Figure S12**). Interestingly, we find that the two most highly abundant repeats in the genome are adjacent to each other and heavily enriched along pericentromeric regions (Figure 4F, **Figure S13**): A 21-bp motif that is found at the center of the centromeric region at most chromosomes, and an unrelated 99-bp repeat motif that is heavily AT-rich, and has characteristics described for other centromeric repeats (Figure 4D, E). Specifically, the 99-bp motif shows a 10-bp periodicity of A and/or T di-and tri-nucleotides, similar to centromere repeats found in diverse species, including *D. melanogaster* or in the legume *Astragalus sinicus* [38,39]. A single turn of the DNA double helix is approximately 10-bp, and sequences with 10-bp periodicity in AA, TT or AT di-nucleotides favor wrapping of nucleosomes by reducing the bending energy of wrapping [40,41]; thus, the 10-bp periodicity of AA, TT or AT di-nucleotides presumably helps to stabilize centromeric nucleosomes that may be under tension during anaphase. Pericentromeric regions are heterochromatic, and we see strong enrichment of H3K9me3 along the pericentromere (Figure 3, 4F). However, centromere repeats are partially occupied by a special centromeric variant of histone H3 (cenH3), which forms specialized nucleosomes that wrap centromeric DNA [42], and we would thus expect less H3K9me3 enrichment at sequences that partly replace the canonical H3 histone with cenH3. Indeed, H3K9me3 enrichment is reduced at the 21-bp motif, and the 99-bp motif, relative to other pericentromeric regions (Figure 4F, **Figure S13**). Thus, the genomic distribution of the 21-bp and 99-bp motif, their structural features and epigenetic modifications strongly suggest that they represent the functional centromere in *D. miranda*.

### Repeat islands along euchromatic chromosome arms

In addition to the repetitive pericentromeres, our assembly also contains two large heterochromatic islands along the two autosomal arms (^~^800 kb on ch2 and 1.5 Mb on ch4; Figure 3). These heterochromatic islands and their positions are supported by *in situ* hybridization data (Figure 1C). Intriguingly, while the repeat density is increased in these islands (especially on ch2), gene density is similar to other euchromatic regions. In *D. melanogaster*, repeat-rich heterochromatic regions appear to be absent along the major chromosome arms, and it will be of great interest to understand the functional significance and phylogenetic distribution of these heterochromatic islands.

### Assembly of the Y and neo-Y chromosome of *D. miranda*

The presence of its recently formed neo-sex chromosomes have established *D. miranda* as an important model system [13,15,19]. Yet, the assembly of both the neo-X and neo-Y proved particularly challenging to short read technology, and our previous attempt to create a contiguous Y/neo-Y chromosome assembly failed [19]. In contrast, our current assembly contains most of the Y chromosome in one large scaffold (101.5 Mb, see Figure 3, 5). Intriguingly, the neo-Y assembly is about 3-times the size of the neo-X assembly (**Table S7**, Figure 5); thus, analysis of neo-Y sequences based on neo-X alignments clearly misses the majority of the changes that occurred between the neo-sex chromosomes.

**Figure 5.**
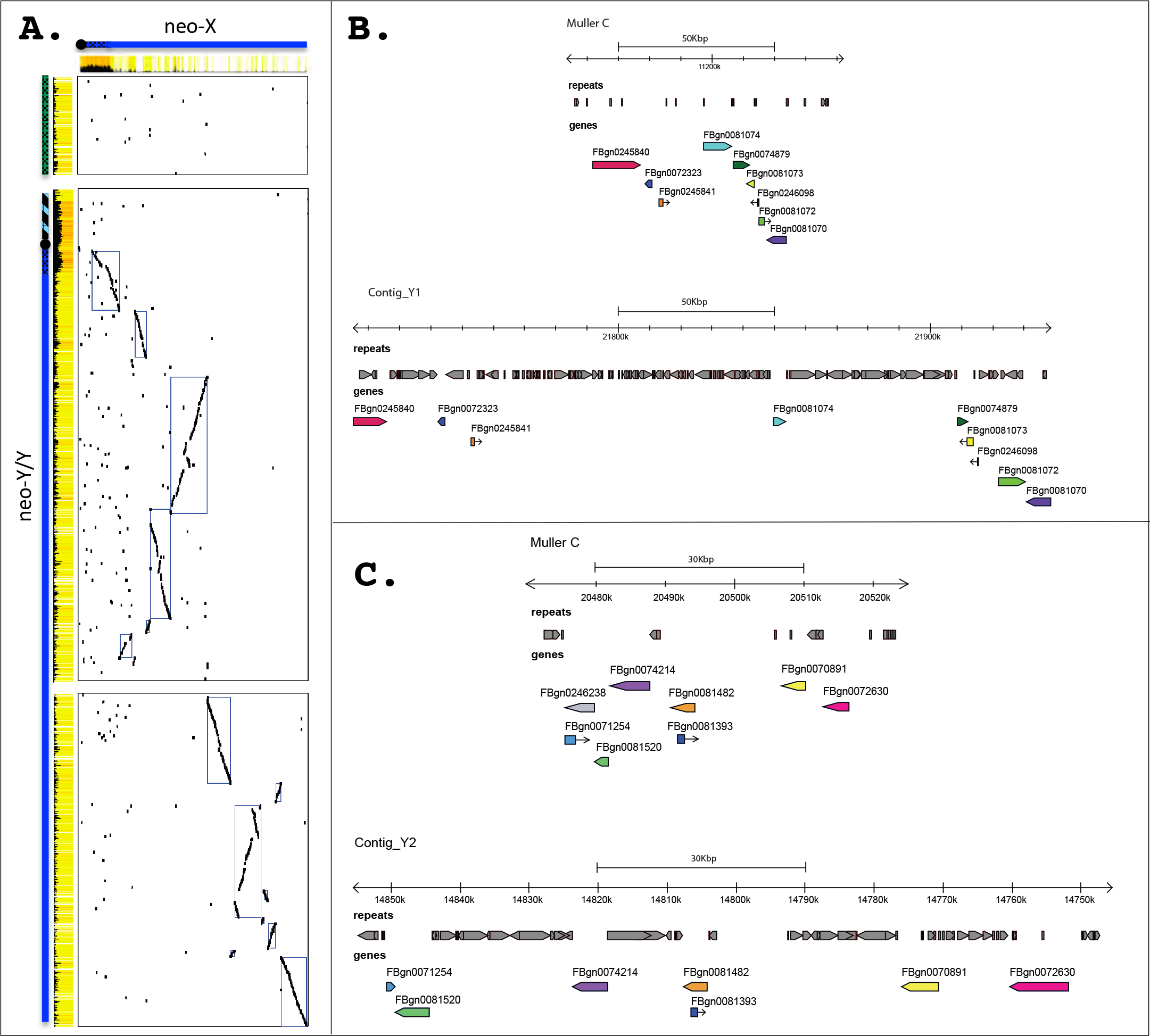
Neo-sex chromosome homology. **A.** Global neo-sex chromosome alignments show large homologous blocks between the neo-sex chromosomes along the long arm of the neo-Y. **B** and **C.** Zoom-in of selected homologous regions along the neo-sex chromosomes. Note the dramatic repeat accumulation at both intergenic and gene regions on the neo-Y, greatly increasing its size.

Sequence analysis of BAC clones confirms that our neo-X and neo-Y assembly is of high quality. In particular, 28 BAC clones fully map to the neo-X chromosome and 92 map to the neo-Y/Y chromosome; only 3 BACs in highly repetitive sequences on the neo-Y map to two different regions (and may either indicate a misassembly or a recombinant BAC clone; **Table S5, S6**; Figure 2D). Thus, our assembly approach allowed us to recover a highly contiguous Y/neo-Y sequence. Inspection of BAC sequences from homologous neo-X and neo-Y regions confirmed the specificity of our neo-X and neo-Y linked assembly. That is, we found little cross-mapping between BAC clone sequences derived from the neo-X chromosome and its former homolog, the neo-Y, and vice versa, confirming the lack of chimeric assemblies (**Figure S14**). Also, comparisons of homologous regions covered by BAC clones confirmed that neo-Y sequences contained roughly 3-times more DNA than their homologous segments on the neo-X, supporting the global size differences in chromosome assemblies that we observe. Thus, rather than shrinking - the fate that is typically ascribed to Y chromosomes - we find that early Y chromosome evolution instead is characterized by a massive global DNA gain.

Genome-wide alignments between the neo-X and neo-Y chromosome support a global increase in size of the neo-Y chromosome at intronic and intergenic regions, mainly driven by the accumulation of repetitive elements (Figure 3, 5, **Figure S15**). We assembled 110.5 Mb of Y/neo-Y linked sequence, and 81.5 Mb are derived from repetitive elements (compared to 5.3 Mb on the neo-X). TEs are uniformly enriched along the neo-Y chromosome (Figure 3, **5**), and a main contributor to its dramatically increased genome size. Transposons show a highly nested structure on the neo-Y, with TE copies being disrupted by the insertion of (fragments of) other transposable elements (Figure 5B, C), making the exact delineation of TE copies challenging. The most abundant repeat on the neo-Y is the ISY element [43], a helitron transposon that is inserted ^~^22,000 times on the neo-Y/Y chromosome, and occupies more than 16 Mb on the neo-Y (Figure 3, **Figure S16**). In contrast, we only find ^~^1,500 copies on its former homolog, the neo-X chromosome (less than 1Mb). The second most common repeat class that has amplified on the neo-Y are gypsy transposable elements; we find roughly ^~^14,300 insertions on the neo-Y (15 Mb), and less than 1 Mb on the neo-X (^~^800 insertions).

### Reconstruction of chromosomal events leading to sex chromosome turn-over

In *D. miranda*, novel sex chromosomes were created recently at two different time points by chromosomal fusions (Figure 1D). In an ancestor of *D. miranda*, about 15MY ago, a new X-linked arm (referred to as chromosome XR) arose by the fusion of an autosome (Muller element D) to the ancestral X chromosome (element A, referred to as chromosome XL in *D. miranda)*. The fusion of Muller element A and D left behind an unfused element D in males (which we refer to as YD), and this chromosome co-segregates with the ancestral Y and is transmitted through males only. The lack of recombination in male Drosophila implies that YD is entirely sheltered from recombination, and thus undergoes genome-wide degeneration [12,44]. Indeed, while the fused Muller D that became part of the X chromosome has maintained most of its ancestral genes (we annotate over 2800 genes on XR), previous attempts to recover Y-linked genes in *D. pseudoobscura* have proven difficult. On one hand, single-copy genes located on the ancestral Y chromosome of Drosophila (Y_anc_) were found to be autosomal in *D. pseudoobscura* and its relatives [16], and located on the small dot chromosome (i.e. element F; [17]). It was suggested that the current Y chromosome in *D. pseudoobscura* instead is the remnant of a highly degenerate YD [16], and we previously identified about 30 transcripts from the Y in *D. pseudoobscura*, most of which were found to have their closest paralogs on Muller D [45]. This supports the idea that the current Y of *D. pseudoobscura* is derived from the unfused Muller D, i.e. YD. A more recent chromosomal fusion (about 1.5 MY ago) between YD and element C created another neo-sex chromosome specific to *D. miranda*. This time, the fused element (termed the neo-Y) is male-limited and undergoing degeneration, while the unfused element (the neo-X), is evolving characteristics typical of X chromosomes [19,29].

Our high-quality genome assembly allows us to reconstruct the evolutionary events leading to the independent creation of novel sex chromosomes in the *D. miranda* genome, and the reversal of a former Y chromosome (Y_anc_) to an autosome (Figures 6). Gene content is conserved across chromosomes in Drosophila (referred to as Muller element’s A-F [46]), with Muller element A being the ancestral X chromosome in the genus Drosophila (Figure 1). We used orthology information from either *D. melanogaster* or *D. pseudoobscura* to infer chromosomal re-arrangements in *D. miranda*, and their evolutionary trajectory (Figure 6).

**Figure 6.**
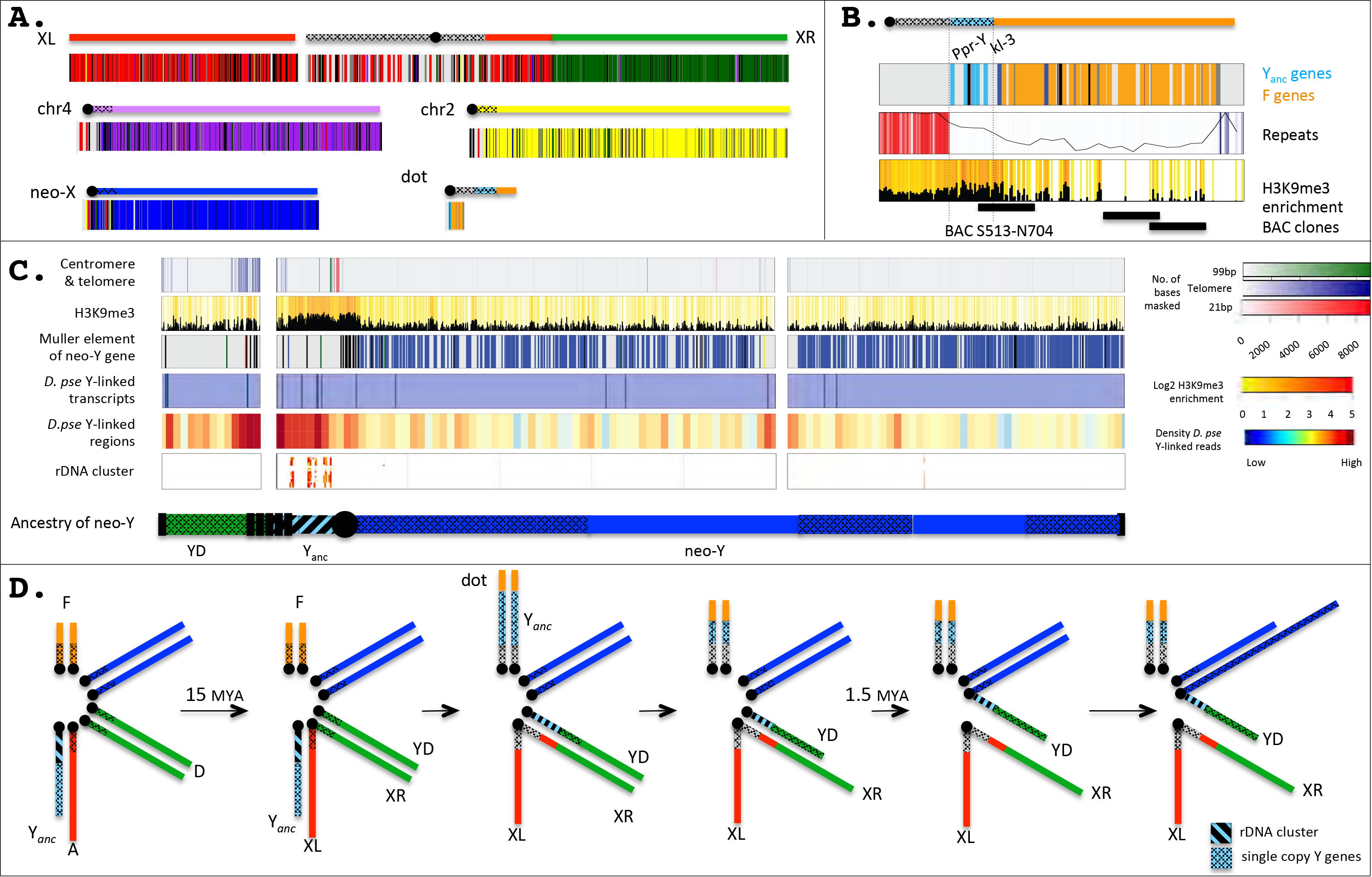
Karyotype evolution in *D. miranda*. **A.** Chromosome arm homology in *D. miranda*. Genes in *D. miranda* are color-coded according to their location in *D. melanogaster* (see Figure 1). **B.** Sequence composition of the *D. miranda* dot. Shown is the origin of dot genes (color coded as in Figure 1), the repeat and H3K9me3 content, as well as the location of sequenced BAC clones. **C.** Origin of the *D. miranda* Y/neo-Y. Shown are the location of centromeric and telomeric repeats, H3K9me3 enrichments, the color coded location of single-copy neo-Y genes, the location of homologous Y-linked genes identified in *D. pseudoobscura*, mapping of Y-derived sequencing reads from *D. pseudoobscura*, and the location of the rDNA genes. The inferred ancestry of the Y/neo-Y chromosome is shown as a cartoon, with the short arm presumably corresponding to the Y chromosome shared with *D. pseudoobscura*, and the long arm representing the neo-Y. **D.** Our genomic analysis allows us to reconstruct sex chromosome evolution in *D. miranda* (see text).

The fusion between element A and D created a metacentric X chromosome in *D. miranda*, and our assembly contains both of these chromosome arms as a single scaffold, including a large pericentromeric block on XR that is highly repeat-rich. Comparison to *D. melanogaster* identifies a pericentric inversion that moved approximately 340 genes from element A onto XR (see Figure 6A). An X chromosome - autosome fusion creates two Y chromosomes in males (i.e. the ancestral Y, and the unfused element D), but *D. miranda* and its relatives only harbor a single Y. The ancestral Y chromosome in Drosophila contains a handful of single-copy genes that have no homologs on the X chromosome [47,48], and the rDNA repeat cluster that is present on both the X and the Y, and used for pairing of the sex chromosomes during male meiosis [49–51]. Our assembly reveals that the gene content of the ancestral Y is split up between two chromosomes: all five ancestral single-copy Y genes in Drosophila (i.e. *kl-2, kl-3*, *ORY*, *PRY* and *PPr-Y)* are located in a single genomic region on element F, adjacent to the centromere (Figure 6B), while the rDNA repeat cluster is found on the Y chromosome of *D. miranda* (Figure 6C). The presence of two Y chromosomes in an ancestor of *D. miranda* may have resulted in an increased frequency of aneuploidy gametes [52], and potential problems in meiosis could have been ameliorated by the fusion and/or translocation of genetic material from Y_anc_ to both element F and YD, and relocation of the rDNA repeat cluster onto YD could have helped to ensure proper segregation between the X and Y chromosome. Indeed, an *in situ* hybridization study suggests that copies of the rDNA loci exist on both the X and Y chromosome in relatives of *D. miranda* that share the element A-D fusion and translocation of single-copy Y_anc_ genes onto element F [17], suggesting that these structural rearrangements co-occurred rapidly before the divergence of this species group.

The Y-derived material on the dot of *D. miranda* amounts to approximately 300 kb, which is substantially smaller than Y chromosomes found in Drosophila [53], suggesting that Y_anc_ presumably lost genetic material after fusing to the dot chromosome. Similar shrinkage of the Y_anc_ was found in its sister species *D. pseudoobscura* [18], which shows an inversion of the Y-derived segment with respect to *D. miranda* (**Figure S17**). The Y_anc_ / element F fusion breakpoint is corroborated independently by a BAC clone spanning the fusion (**Figure S17**), validating our genome assembly in this region. Despite Y_anc_ presumably having lost large amounts of repetitive DNA, we find its repeat content to be elevated, relative to euchromatic regions, and Y_anc_ genes contain higher levels of heterochromatin compared to genes from other chromosomes (**Figure S17**); they also maintained their testis-specific expression pattern in *D. miranda* (**Figure S17**). Thus, despite having become linked to an autosome, single-copy Y genes have retained their ancestral chromatin environment and testis function.

Non-recombining Y chromosomes degenerate within a few million years in Drosophila [19,29], and most ancestral genes on YD were presumably lost before it fused to element C about 1MY ago. We tried to reconstruct the evolutionary history of the Y chromosome in *D. miranda*, by identifying which parts of the Y/neo-Y chromosome were derived from Muller D *vs*. Muller C vs. the original Y_anc_. Our Y/neo-Y chromosome assembly consists of two chromosome arms, spanning the putative centromeric repeats, and the heterochromatic pericentromere (Figure 6C). A dot plot between the neo-X (Muller C) and neo-Y reveals several large blocks of homology on the large Y/neo-Y arm, but none on the shorter arm (Figure 5A). Figure 6C plots the location of single-copy genes along the neo-Y/Y chromosome of *D. miranda*, color-coded by Muller element. We identify many genes from the long arm of the Y/neo-Y, most of which are derived from Muller C; in contrast, only few unique genes exist on the short arm, and their closest homologs are not preferentially located on Muller C (Figure 6C). This suggests that the long arm is derived from the neo-Y, but not the shorter one, which instead may be derived from YD and should thus also be Y-linked in *D. pseudoobscura*. The current genome of *D. pseudoobscura* lacks an assembly of its Y chromosome and repetitive non-functional regions evolve rapidly, which makes identification of YD sequences challenging. We attempted to detect putative YD sequences by identifying reads and scaffolds from the fragmented *D. pseudoobscura* genome that are male-specific (see Methods), and mapping them onto our *D. miranda* Y/neo-Y assembly. Preferential mapping of putative male-specific (i.e. Y-linked) sequences from *D. pseudoobscura* to the short arm of the *D. miranda* Y/neo-Y chromosome assembly supports the notion that the short arm of the Y/neo-Y chromosome corresponds to YD. The rDNA cluster maps adjacent to the centromere on the short arm of the Y chromosome, which suggests that this part is derived from the original Y (i.e. Y_anc_) of Drosophila.

Interestingly, hybridization studies have shown that the Y/neo-Y chromosome of *D. miranda* contains ^~^70 copies of the telomere repeat [33] and displays an intensely labeled internal telomere-repeat block adjacent to the centromere [54]. Indeed, our assembly recovers a large internal block of telomere repeat sequences close to the centromere (Figure 6C), bordering fragments of the Y chromosome of different evolutionary origin (i.e. they are found between fragments derived from Muller D vs. Muller C vs. the original Y_anc_). Telomere repeats within the Y/neo-Y may present the remnants of a ‘telomere-to-telomere’ type chromosomal fusion that created the neo-Y/Y chromosomal arrangements in this species.

**Conclusion:** Here, we create a genome assembly of unprecedented quality and contiguity for the fruit fly *D. miranda*, a species that has served as a model for sex chromosome research. In *D. miranda*, chromosomal fusions at different time points independently created *de novo* sex chromosomes, or led to the reversal of a former Y to an autosome, and our high-quality assembly allows us to reconstruct the evolutionary events creating and dismantling sex chromosomes. Our assembly recovers entire chromosomes and notoriously difficult regions to assemble, including entire centromeres, or large stretches of repetitive sequences, such as the rDNA cluster. All chromosome arms of *D. miranda* - including its Y chromosome - are contained in a single, chromosome-sized scaffold, and in almost all cases, chromosome arms are flanked by telomere sequences on one end, and centromeric repeats on the other. In one instance, we assemble an entire chromosome, and fully sequence through the pericentromeric DNA and the centromere, and recover telomeres on both ends. Our high-quality assembly allows us to infer the centromeric satellite DNA motif in *D. miranda*, which shares no sequence similarity with other centromeres but has characteristics typical of centromeric repeats, including a 10-bp periodicity of AA/TT/AT repeats. Lack of sequence conservation confirms that centromeres turn over quickly [36], and will allow the functional characterization and investigation of centromere biology in this group. For the first time, we also assemble an entire Y chromosome using shot-gun sequencing approaches. In particular, the recovered Y/neo-Y sequence is over 100Mb large, which is over 3-times the size of that of its former homolog, the neo-X or its autosomal ortholog in *D. pseudoobscura*. Thus, rather than shrinking - the fate that is typically ascribed to Y chromosomes - we find that early Y chromosome evolution instead is characterized by a global DNA gain. We show that the *D. miranda* Y chromosome provides a hodgepodge of sequences that have been male-limited for different amounts of time, and display various stages of degeneration. Our new highly improved genome assembly will provide the basis for further evolutionary and functional research on repetitive sequences and the recently formed neo-sex chromosomes of *D. miranda*.

## Methods

### Fly strain

We chose the inbred MSH22 strain for *D. miranda*, which was previously used to generate a BAC library [29], and for genome assembly using short Illumina reads [19].

### PacBio DNA extraction and genome sequencing

We used a mix of MSH22 males and extracted high molecular weight DNA using a QIAGEN Gentra Puregene Tissue Kit (Cat #158667), which produced fragments>100 kbp (measured using pulsed-field gel electrophoresis). DNA was sequenced on the PacBio RS II platform. In total, this produced 28 Gb spanning 2,407,465 filtered subreads with a mean read length of 12,818 bp and an N50 of 17,116 bp (**Table S1**, **Figure S18**).

### BioNano DNA extraction and optical mapping

DNA was extracted from flash frozen male larvae. Purified DNA was embedded in a thin agarose layer and was labeled and counterstained following the IrysPrep Reagent Kit protocol (BioNano Genomics). Samples were then loaded into IrysChips and run on the Irys imaging instrument (BioNano Genomics). This produced 90,977 molecules (molecule length: min 150,000, median 191,400 and max 1,957,000 and N50 of 209,014; **Table S9**; **Figure S19**). The IrysView (BioNano Genomics) software package was used to produce single-molecule maps and *de novo* assemble maps into a genome map (**Table S4**). The BioNano assembly has 401 contigs with an N50 of 0.5 Mb and assembled length of ^~^178Mb. HybridScaffold was then used to produce hybrid maps from the BioNano contigs and the genomic scaffolds from our scaffolded PacBio assembly, and IrysView was used to visualize alignments of the BioNano contigs and genomic scaffolds to the hybrid ones. **Table S4** shows coverage of hybrid scaffolds by BioNano contigs and NGS contigs (genomic scaffolds).

### PacBio assembly

40x error corrected reads were used to build an initial PacBio assembly using the Falcon assembler [22]. 28Gb long reads (NR50 = 17116 bp; NR50 is the read length such that 50% of the total sequence is contained within reads of this length or longer) were assembled using Falcon assembler (v1.7.5) [22] running on Sun Grid Engine in parallel mode. For assembly, reads longer than 10Kb and 17Kb were used as seed reads for initial mapping and pre-assembly. The options for read correction, overlap filtering, and consensus building were provided in the config file as follows: pa_HPCdaligner_option = -v -dal128 -t16 -e.70 -l1000 -s1000; ovlp_HPCdaligner_option = -v -dal128 -t32 -h60 -e.96 -l500 -s1000; pa_DBsplit_option = -x500 -s400; ovlp_DBsplit_option = -x500 -s400; falcon_sense_option = -output_multi -minjdt 0.70 -min_cov 4 -max_n_read 200 --n_core 6; overlap_filtering_setting = -max_diff 30 -max_cov 60 -min_cov 5 --n_core 24. This assembly had 629 scaffolds and a total assembled length of 274,803,116 bp with an N50 value equal to 2,188,952 bp. We polished this assembly using the software Quiver [55], followed by the software Pilon [56] which resulted in an assembly with 625 scaffolds, with an N50 value of 2,232,625 bp and total assembled length equal to 271,223,447 bp. We also produced a second PacBio assembly using Canu [23] with default parameters. This assembly consisted of 521 scaffolds and a total assembled length of 296,012,170 bp, with an N50 value of 3,884,273 bp. These two assemblies were the merged using Quickmerge [24], with default parameters. The resulting merged assembly was then merged a second time to the finished Falcon assembly, producing a Quickmerged assembly consisting of 271 scaffolds and total length equal to 295,213,648 bp and an N50 value of 5,177,776 bp.

### Hi-C libraries

Hi-C libraries were created from sexed male and female 3^rd^ instar larvae of MSH22 following [57]. Briefly, chromatin was isolated from male and female 3rd instar larvae of *D. miranda*, fixed using formaldehyde at a final concentration of 1%, and then digested overnight with HindIII and HpyCh4IV. The resulting sticky ends were then filled in and marked with biotin-14-dCTP, and dilute blunt end ligation was performed for 4 hours at room temperature. Cross-links were then reversed, and DNA was purified and sheared using a Covaris instrument LE220. Following size selection, biotinylated fragments were enriched using streptavidin beads, and the resulting fragments were subjected to standard library preparation following the Illumina TruSeq protocol. For females, 38.4 and 194.5 million 100-bp read pairs were produced for the HpychIV and HindIII libraries, respectively. For males, 28.0 and 179.2 million pairs were produced.

### Hi-C-based proximity guided assembly (PG)

We mapped Illumina male and female genomic paired-end reads and classified contigs as autosomal, X-linked or Y-linked based on genomic coverage. We created two pools of contigs: autosomes or X-linked, and Y-linked, and scaffolded them separately. We used Juicer [58] to align female Hi-C reads to the autosomal/X-linked scaffolds and also to align a subset of male Hi-C reads (that did not map to autosomes) to the Y-linked scaffolds. There were 22,168,695 Hi-C contacts - 2,921,250 interchromosomal and 19,247,445 intrachromosomal contacts for the autosomal/X linked scaffolds. For the Y linked scaffolds, there were 795,487 Hi-C contacts, including 173,147 interchromosomal and 622,340 intrachromosomal contacts. The output alignment files from Juicer were then used to scaffold the genome using 3D-DNA [59]. Using a custom Perl script, we then scaffolded the Pac-Bio assembly fasta based on the 3D-DNA output suffixed.asm, which contains information about the positions and orientations of contigs; scaffolded contigs are gapped by 50 Ns. With the Hi-C scaffolded assembly, we then realign the Hi-C reads using bwa mem [60] single-end mode on default settings. The resulting *sam* files were then used to generate a genome-wide Hi-C interaction matrix using the program Homer [61]. For visualization, we plotted the interaction matrix as a heatmap in R, with demarcations of the Pac-Bio contigs and Hi-C scaffolds. Iteratively, we visually examine the heatmap to identify possible anomalies as scaffolding errors and manually curate the.asm file output to improve the heatmap. At each stage of the assembly process, genome completeness was assessed using BUSCO (v 3.01) [28], using the arthropod database (odb9).

### BAC clone DNA isolation and sequencing

Bacteria were cultured in Terrific Broth with 25 μg/mL chloramphenicol. Overnight cultures (500 μl) were inoculated with starter cultures grown from glycerol stocks, covered with AreaSeal films, and incubated at 37°C with shaking for 12–14 hours. Overnight cultures were pelleted by centrifugation, re-suspended in 60 μl [Tris-HCl (50 mM, pH 8) and EDTA (50 mM)], and lysed by adding 120 μl [NaOH (200 mM) and SLS (1%)]. Cells were incubated at room temperature for five minutes, 270 μl [KOAc (5 M, pH 5)] was added and chilled on ice for 10 minutes, and then centrifuged for 1 hour. DNA was precipitated with isopropanol, washed with 70% and 80% ethanol and eluted in Qiagen EB (50 μl). Nextera libraries were prepared from the BAC DNA, following Illumina’s protocol with the following modifications: reaction volumes were scaled to 1 μl input BAC DNA (@ 1-3 ng/μl), and SPRI bead cleanup steps after tagmentation and PCR amplification were skipped. Barcoded libraries were pooled, and a 2-sided Ampure XP size-selection removed fragments <200bp and minimized fragments >800bp. The pooled libraries were sequenced on a HiSeq 4000 with 100 bp paired-end reads.

### BAC clone mapping

For each BAC clone, Nextera reads were first adapter trimmed using cutadapt (http://code.google.com/p7cutadapt/) and filtered to remove concordantly mapping read pairs from pTARBAC-2.1 and *E. coli* DH10B using Bowtie2 [62]and SAMtools [63]. The remaining trimmed, filtered reads were mapped to our *D. miranda* assembly using bwa [60]. The BAC’s location was determined by filtering regions of high coverage (at least 50X mean) and significant length (at least 20-kb). First, regions with average coverage of at least 50X were extracted, and any regions within 250-kb of each other were merged using BEDtools [64]. When this resulted in a merged region longer than 250-kb, the merging step was repeated on this long region using a maximum distance of 5-kb. If only one region remained, this was defined as the putative BAC location. If multiple regions were found, they were ranked by average coverage, and any region with less than half the average coverage of the region with the highest average coverage was considered cross contamination. Finally, regions less than 20-kb long were removed.

To confirm that reads mapping to these BAC locations included both edges of the BAC insert, we found discordantly mapping read pairs with one read mapping to the vector and its mate mapping to our assembly. Filtered reads were mapped to pTARBAC-2.1 using bwa [60], and discordantly mapping reads from either end were filtered from the.sam file, keeping "start" and "end" reads separated. (Reads mapping to a region within 4000-bp of the vector’s start position were considered “start” reads, and reads mapping within 4000-bp of the vector’s end position were considered "end" reads.) The mates of these start/end reads were extracted, merged and counted using BEDtools [64], filtered to find edge read pileups within 10-kb of the putative BAC edges. To confirm that these edge reads are at either end of each BAC location, IGV snapshots with three tracks (all mapped reads, "start" reads, and "end" reads) were reviewed manually.

To confirm that our assembly of the neo-X and neo-Y were highly specific and accurate, the genomic region on the neo-sex chromosome from which a specific BAC clone was derived was masked using BEDtools [64], and the BAC clone reads were mapped back to this masked assembly, and then filtered and merged as described above. Regions of primary and secondary mapping were reviewed using IGV to confirm that little cross mapping occurs in our assembly; after masking and re-mapping, we found significant mapping to homologous regions of the its homologous neo-sex chromosome, but mapped reads typically contained many SNP’s and many gapped regions (**Figure S14**).

### Conflict resolution

To identify large-scale, erroneously duplicated regions, we took advantage of the fact that when reads are mapped equally well to multiple regions, they are randomly assigned to one of the regions; we mapped illumina reads to the assembly twice and identified >100kb regions where roughly half of the reads map to another region in the two mappings (see **Figure S4**). For erroneous duplications and mis-scaffolded contigs in the Pac-Bio assembly identified, we used IGV to visualize the quality of Illumina reads mapping, in order to determine the precise coordinates to modify our assembly (**Table S2**). For erroneous duplications, we identified the position in which Illumina reads are no longer uniquely mapping around the duplicated areas; one of the two duplications are then removed. Misscaffolded contigs are typically caused by misassembly around repetitive elements, therefore we also relied on visual inspection of non-uniquely mapping reads to separate contigs.

### Structural Variant Calling for Quality Control

For the previously published genome assembly and the various intermediate assemblies produced here during generating the current version, we estimated quality statistics using the variant caller LUMPY [65]. To do this, we first aligned reads from two separate male Illumina libraries (with 626bp and 915bp insert sizes, respectively) to our current assembly and its intermediates using SpeedSeq, which does a BWA-MEM alignment and produces discordant and split reads bam files. We ran the software lumpyexpress [65] using these bam files which produced a vcf file with several categories of structural variants: BND =trans-contig associations, DEL = deletions, DUP= Duplications, INV= Inversions. High numbers of these variants are indicative of potential assembly errors and provide a meaningful way to assess assembly quality.

Repeat Annotation and masking. For repeat masking the genome, we annotated repeats using REPdenovo (downloaded November 7, 2016; [25]) and RepeatModeler version 1.0.5 [26]. We ran REPdenovo on raw sequencing reads using the parameters MINREPEATFREQ 3, RANGEASMFREQDEC 2, RANGEASMFREQGAP 0.8, KMIN 30, KMAX 50, KINC 10, KDFT 30, GENOMELENGTH 176000000, ASMNODELENGTHOFFSET-1, MINCONTIGLENGTH 100, ISDUPLICATEREPEATS 0.85, COVDIFFCUTOFF 0.5, MINSUPPORTPAIRS 20, MINFULLYMAPRATIO 0.2, TRSIMILARITY 0.85, and RMCTNCUTOFF 0.9. We ran RepeatModeler with the default parameters.

We used tblastn (https://www.ncbi.nlm.nih.gov/BLAST/) with the parameters -evalue 1e-6, -numalignments 1, and -numdescriptions 1 to blast translated *D. pseudoobscura* genes (release 3.04) from FlyBase [66] to both (REPdenovo and RepeatModeler) repeat libraries. We eliminated any repeats with blast hits to *D. pseudoobscura* genes. After filtering, our REPdenovo repeat annotation had 999 repeats totaling 964,435 base pairs.

We also made a REPdenovo annotation using a subset of female reads, for which we also filtered out repeats blasting to *D. pseudoobscura* genes. This annotation had 716 repeats totaling 544,702 base pairs. We used RepeatMasker version 4.0.6 [26] and blastn (https://www.ncbi.nlm.nih.gov/BLAST/) with the parameters -evalue 1e-6, -numalignments 1, and -numdescriptions 1 to blast this annotation to the Repbase Drosophila repeat annotation (downloaded March 22, 2016 from http://www.girinst.org) in order to classify repeats from this annotation. Our RepeatModeler repeat annotation had 1,009 repeats totaling 1,290,513 base pairs. Of the 1,009 repeats, 103 were annotated as DNA transposons, 145 as LINEs, 365 as LTR transposons, 42 as Helitrons, and 1 as a SINE. We concatenated our filtered REPdenovo and RepeatModeler repeat annotations to repeat mask the genome with RepeatMasker [67].

### Gene annotation using Maker

To run Maker [27], we first build transcriptome assemblies. RNA-seq reads from several adult tissues (male and female heads, male and female gonads, male accessory gland, female spermatheca, male and female carcass, male and female whole body and whole male and female 3rd instar larvae; see **Table S10**) were aligned to the genome assembly using HiSat2 [68] using default parameters and the parameter -dta needed for downstream transcriptome assembly. The alignment produced by HiSat2 was then used to build a transcriptome assembly using the software StringTie [69] with default parameters, which produced a transcript file in gtf format. Fasta sequences of the transcripts were then extracted using gffread to be used with Maker. The genome was repeat masked using RepeatMasker and our *de novo* repeat library as well as the Repbase (http://www.girinst.org/) annotation.

We ran three rounds of Maker [27] to iteratively annotate the genome. For the first Maker run, we used annotated protein sequences from flybase for *D. melanogaster* and *D. pseudoobscura*, as well as the *de novo* assembled *D. miranda* transcripts and the genes predictors SNAP [70] and Augustus [71] to guide the annotation. We used the SNAP *D. melanogaster* hmm and the Augustus fly model, with the parameters est2genome and protein2genome set to 1 in order to allow Maker to create gene models from the protein and transcript alignments. Before running Maker a second time, we first trained SNAP using the results of the previous Maker run and set the est2genome and protein2genome parameters to 0. We then used our new hmm file and the Augustus fly model to annotate the genome. The 3rd iteration was done similarly to the second one, by training SNAP on the results of the previous Maker run. This procedure resulted in a total of 17,745 annotated genes. The repeat and gene densities were plotted for the major chromosomal arms and scaffolds using the software DensityMap [72].

### Tandem repeat identification and quantification

We used Tandem Repeat Finder (TRF) [37] on recommended settings to identify tandemly repeating motifs across the assembly. To identify variants or multimers of the same motif, the identified motifs are then blasted pairwise to themselves. Those that are over 90% identical for over 90% of the length are grouped together and collapsed into the same motif. Satellites abundances were parsed from the TRF output and RepeatMasker output using the identified motifs as the repeat library.

### Identifying telomeric repeats

Telomeric protein sequences for *D. pseudoobscura* and *D. persimilis* from [32] were aligned to the *de novo* repeat library using BLAST. Hits with a score greater than 50 and percent identity greater than 75 were classified as telomeric and *Repeatmasker* was used to identify their genomic locations. A heatmap showing the number of bases masked in 10Kb windows was then plotted along the genome using R.

### Identifying orthologous proteins and whole-genome alignments

We identified orthologous proteins by aligning *D. pseudoobscura* proteins to our list of *de novo* annotated *D. miranda* proteins using BLAST and BLAT. For 16,378 of the total 17,745 genes in our annotation we were able to reliably identify orthologs in the *D. pseudoobscura* annotation. We used blastp to align protein sequences of the remaining 1,367 genes to annotated *D. melanogaster* proteins and were able to identify *D. melanogaster* orthologs for 285 of these 1,367 genes. Thus, we were unable to identify orthologs for 1,082 genes in both the *D. pseudoobscura* and the *D. melanogaster* genome. Whole genome alignments were performed using Nucmer (from the MUMmer package [73]) and dotplots were produced using mummerplot, symap42 [74] or YASS [75].

### Identifying *D. pseudoobscura* Y-linked reads

Scaffolds from a male-only *D. pseudoobscura* assembly were aligned to a female only *D.pseudoobscura* assembly using Nucmer (from the MUMmer package [73] to identify putative Y-linked scaffolds. Male Illumina reads were then aligned to these scaffolds using bowtie2 [62] and unaligned reads were discarded. The aligned reads were then mapped to the female genome to further enrich for only male-specific reads. These reads were then mapped to the D. miranda Y/neo-Y linked scaffolds and coverage was calculated in 10Kb non overlapping windows. The density of non-zero coverage windows was plotted along the three largest Y scaffolds.

## Acknowledgements

Funded by NIH grants (R01GM076007, GM101255 and R01GM093182) to DB. We thank JJ Emerson, Mahul Chakraborty, Ryan Bracewell, Emily Brown and Alison Nguyen for technical assistance.

## Author contributions

SM and DB conceived the study, SM, KW, MN and LG collected and analyzed the data and SM and DB wrote the manuscript with input from all authors.

